# Protein appetite drives macronutrient-related differences in ventral tegmental area neural activity

**DOI:** 10.1101/542340

**Authors:** Giulia Chiacchierini, Fabien Naneix, Kate Zara Peters, John Apergis-Schoute, Eelke Mirthe Simone Snoeren, James Edgar McCutcheon

**Affiliations:** Dept. of Neuroscience, Psychology & Behaviour, University of Leicester, University Road, Leicester, LE1 9HN, United Kingdom; The Rowett Institute, University of Aberdeen, Foresterhill, Aberdeen AB25 2ZD, United Kingdom; Dept. of Psychology, UiT The Arctic University of Norway, Huginbakken 32, 9037 Tromsø, Norway

## Abstract

Control of protein intake is essential for numerous biological processes as several amino acids cannot be synthesized *de novo*, however, its neurobiological substrates are still poorly understood. In the present study, we combined in vivo fiber photometry with nutrient-conditioned flavor in a rat model of protein appetite to record neuronal activity in the ventral tegmental area (VTA), a central brain region for the control of food-related processes. In adult male rats, protein restriction increased preference for casein (protein) over maltodextrin (carbohydrate). Moreover, protein consumption was associated with a greater VTA response relative to carbohydrate. After initial nutrient preference, a switch from a normal balanced diet to protein restriction induced rapid development of protein preference but required extensive exposure to macronutrient solutions to induce greater VTA responses to casein. Furthermore, prior protein restriction induced long-lasting food preference and VTA responses. This study reveals that VTA circuits are involved in protein appetite in times of need, a crucial process for all animals to acquire an adequate amount of protein in their diet.

**Significance Statement:** Acquiring insufficient protein in one’s diet has severe consequences for health and ultimately will lead to death. In addition, a low level of dietary protein has been proposed as a driver of obesity as it can leverage up intake of fat and carbohydrate. However, much remains unknown about the role of the brain in ensuring adequate intake of protein. Here, we show that in a state of protein restriction a key node in brain reward circuitry, the ventral tegmental area, is activated more strongly during consumption of protein than carbohydrate. Moreover, although rats’ behavior changed to reflect new protein status, patterns of neural activity were more persistent and only loosely linked to protein status.

## Introduction

Ensuring appropriate intake of the three main macronutrients (carbohydrate, fat, protein) is a compelling problem for survival of all animals, including humans. Of the three macronutrients, protein intake is thought to be the most tightly regulated, as many amino acids cannot be synthesized *de novo* (Berthoud et al., 2012). Concordantly, many species, including invertebrates (Mayntz et al., 2005) and mammals (Theall et al., 1984), adjust their behavior to ensure adequate intake of dietary protein. In humans, inadequate protein levels in diet may contribute to obesity, by leveraging up the amount of calories consumed from fats and sugar (Simpson and Raubenheimer, 2005; Hall, 2019; Raubenheimer and Simpson, 2019). Recently, we developed a rodent model of protein appetite in which animals rodents maintained on a protein-restricted diet developed a strong preference for a protein-rich solution, relative to a carbohydrate-rich solution (Murphy et al., 2018; see also Hill et al., 2019), indicating that animals can specifically direct feeding and food-seeking behavior towards protein sources in times of need. However, the neural mechanisms by which diets that are low in protein might shift behavior are not understood.

The ventral tegmental area (VTA) and its projections play a central role in food-seeking behaviors, food preference, and in the motivation to eat (Ikemoto and Panksepp, 1999; Berridge, 2007; Bromberg-Martin et al., 2010). VTA neurons are sensitive to numerous food-related signals, including ingestive and post-ingestive processes (de Araujo et al., 2008; Domingos et al., 2011; Beeler et al., 2012; Ferreira et al., 2012; McCutcheon et al., 2012a; Alhadeff et al., 2019), and peripheral hormones (Di Chiara and Abizaid, 2009; Mebel et al., 2012; Mietlicki-Baase et al., 2013, 2014; Cone et al., 2014), allowing the formation of future food preferences (Sclafani et al., 2011). Despite abundant data on the involvement of VTA activity in mediating responses to fat- or carbohydrate-containing food, the role of this region in regulation of protein appetite is still unexplored.

Here, we use *in vivo* fiber photometry to record the activity of VTA neurons during consumption of isocaloric protein- and carbohydrate-containing solutions in an animal model of protein preference (Murphy et al., 2018; Naneix et al., 2019, 2020). We find that, in protein-restricted animals, protein consumption is associated with elevated neural activation, relative to carbohydrate consumption. We then show that when physiological state is reversed behavioral protein preference shifts to reflect the new state more rapidly than neural activity in the VTA.

## Materials and Methods

### Subjects

Adult male Sprague Dawley rats (Charles River Laboratories, n=15) weighing 250-300g on arrival were used. Rats were housed in pairs in individually ventilated cages (46.2 x 40.3 x 40.4 cm), in a temperature (21 ± 2°C) and humidity (40-50%) controlled environment with a 12 h light/dark cycle (lights on at 7:00 AM) and with water and food available *ab libitum*. All testing occurred in the light phase. Data are not reported for seven rats due to poor or non-existent photometry signal resulting from lack of viral expression, misplacement of fiber, or poor connection between patch cable and ferrule. Two rats were removed from the study due to aggressive behavior in the week following the initial dietary manipulation, which led to them being singly housed, rather than in pairs. Procedures were performed in accordance with the Animals (Scientific Procedures) Act 1986 and carried out under Project License 70/8069 / PFACC16E2.

### Virus Injection and Fiber Implantation

For fiber photometry recording, rats received a unilateral injection of a GCaMP6s expressing virus in the VTA and were implanted with fiber optic cannulas targeting the injection site (Fig. 1A). One-two weeks after their arrival, rats were anesthetized with isoflurane (5% induction, 2-3% maintenance) and mounted in a stereotaxic frame (David Kopf Instruments) in a flat skull position. The scalp was shaved, cleaned with chlorhexidine and locally anaesthetized with bupivacaine (150 µl, s.c.). Rats also received i.p. injection of non-steroidal anti-inflammatory meloxicam (1 mg/kg). Core body temperature, oxygen saturation and heart rate were monitored throughout the surgery. A hole was drilled above the VTA at the following coordinates: AP -5.8 mm, ML +0.7 mm relative to Bregma (Paxinos and Watson, 1998). A 10 µl Hamilton syringe placed in a motorized syringe pump (Harvard Apparatus Pump 11 Elite) was loaded with the GCaMP6s virus (AAV9.Syn1.GCaMP6s.WPRE.SV40, ≈1.9x10^13^ GC/ml, Penn Vector Core; RRID: Addgene_100843) and was slowly lowered into VTA (DV -8.1 mm relative to brain surface). 1 μl of virus was delivered over 10 minutes (100 nl/min) and the syringe was left in place for 5 additional minutes before being slowly removed. An optic fiber cannula (ThorLabs CFM14L10, 400 μm, 0.39 NA, 10 mm length) was implanted at the same coordinates, 0.1 mm above the injection site (DV -8.0 mm relative to brain surface). The cannula was secured in place by dental cement (C&B Supabond followed by regular dental acrylic, Prestige Dental) overlaying 4 small skull-screws. Rats were housed in pairs immediately for recovery. Rats were allowed at least 4 weeks to recover before the start of behavioral testing to allow ample time for virus expression.

**Figure 1.**
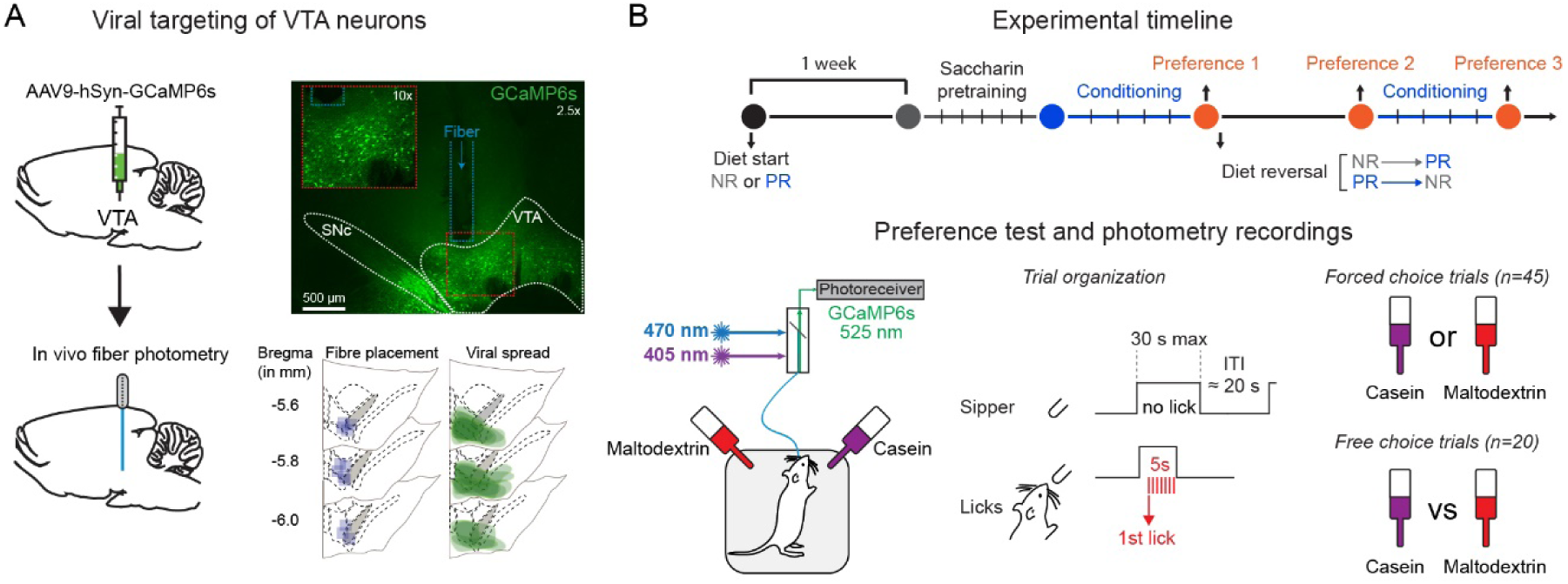
Experimental procedures and timeline. ***A.*** Schematic showing targeting of ventral tegmental area (VTA) by GCaMP6s and implantation of optic fiber (left). Expression of virus in VTA and fiber track are shown in photomicrograph (top right) and location of expression and fiber placements are shown for all rats (bottom right). ***B.*** Schematic showing experimental timeline (top), fiber photometry set-up (bottom left), and trial structure of preference tests (bottom right).

### Diets

All rats were initially maintained on standard laboratory chow diet (EURodent Diet 5LF2, LabDiet) containing 14% protein. Four weeks after surgery, eight of the rats were randomly assigned to the protein-restricted diet condition (PR). For these rats, standard chow was switched to a modified AIN-93G diet containing 5% protein from casein (#D15100602, Research Diets; Murphy et al., 2018). Remaining rats were maintained under standard laboratory chow diet (non-restricted group, NR). Behavioral testing started 1 week following protein restriction.

### Flavor Conditioning and Casein Preference tests

Animals were trained in two identical conditioning chambers (30.5 x 24.1 x 21.0 cm; Med Associates), each located inside a sound- and light-attenuated aluminum outer chamber (1200 x 700 x 700 cm). Each conditioning chamber was equipped with a house light located on the left wall, 2 retractable sippers located on the right wall and 2 light cues located above each sipper hole. Each bottle placed on a retractable sipper was connected to a contact lickometer (Med Associates) used to measure intake of flavored solution. The house light was turned on at the beginning of each daily session and turned off at the end of it. Conditioning chamber apparatus was controlled via a computer running Med- PC IV Software Suite (Med Associates). Sessions were video recorded at either 5 Hz or 10 Hz using a webcam (Microsoft LifeCam) that interfaced with fiber photometry software.

Initially, all rats were pretrained with 2 bottles containing 0.2% sodium saccharin (Sigma). First, rats had continuous access to both bottles in the chambers until they reached >1000 licks during the daily 60 min session (1-3 days). Then, each saccharin bottle was presented individually in a pseudorandom order (inter-trial interval 10-30 s, mean 20 s) during 45 trials On each trial, if no licks were made, then sippers remained available for 30 s. However, once a lick was made, sippers remained extended for 5 s before retraction (Fig. 1B). This protocol trained rats over a small number of sessions to approach and drink from sippers when available. Coincident with sipper activation, the cue light located above the sipper hole was turned on and remained on until the sipper was retracted. Sippers took approximately 2 s from activation until the rat could reach them to drink. Rats were trained with 0.2% saccharin (sodium salt hydrate, Sigma #S1002) in both bottles until they reached the criteria of >1000 licks across the session. Following saccharin pre-training, during the next 4 days, all rats were trained to associate a specific flavored solution (0.05% cherry or grape Kool-Aid with 0.2% saccharin) with a different nutrient in daily sessions lasting a maximum of 60 min. (**Conditioning sessions**). During conditioning sessions, only one bottle was available and was presented during 45 individual trials, as described above. Bottles were filled with either protein-containing solution (4% casein, sodium salt from bovine milk, Sigma #C8654; 0.21% L-methionine, Sigma #M9625; 0.2% saccharin; 0.05% flavored Kool-Aid) or isocaloric carbohydrate-containing solution (4% maltodextrin, Sigma# 419672; 0.2% saccharin; 0.05% flavored Kool-Aid), as previously described (Murphy et al., 2018). Bottle positions, presentation order, and flavor-macronutrient associations were counterbalanced between rats. Bottle position was alternated between days.

Twenty-four hours after the last conditioning session, rats received a first preference test (**Pref test 1**). Both casein and maltodextrin-flavored solutions were available during the test. The test started with 45 trials during which each bottle was presented in pseudorandom order (**Forced choice trials**; 20 sec variable inter-trial interval). These trials were followed by 20 presentations of the two bottles simultaneously (**Free choice trials**).

Immediately after Preference test 1, diet conditions were switched between experimental groups. Non-restricted rats were now given protein restricted diet (**NR**→**PR**) while protein restricted rats were given standard chow diet (**PR**→**NR**). Seven days after the diet switch, a second preference test was conducted (**Pref test 2**). This test was followed by 4 days of additional conditioning sessions, as described above, before a final preference test (**Pref test 3**).

### Fiber Photometry Recordings

To assess the activity of VTA neurons during the consumption of differently-flavored macronutrient solutions, the ‘bulk’ fluorescence signal generated by GCaMP6s expressing cells was recorded using fiber photometry (Fig. 1; Gunaydin et al., 2014; Lerner et al., 2015). Signal processing and acquisition hardware (RZ5P; Tucker Davis Technologies) was used to control two light sources: a 470 nm LED (ThorLabs, M470F3) modulated at 211 Hz and a 405 nm LED (ThorLabs, M405F1) modulated at 539 Hz. A fluorescence minicube (Doric Lenses) combined both wavelengths, which were transmitted through an optical patch cable to the rats’ optic cannula implant. LED power was set at 30-60 µW. Emitted light was delivered through the same patch cable back to the minicube where it was filtered for GFP emission wavelength (525 nm) and sent to a photoreceiver (#2151 Femtowatt Silicon Photoreceiver, DC-750 Hz; Newport). Demodulation of the two light sources allowed dissociation of calcium-dependent GCaMP6s signals (470 nm) and calcium-independent changes resulting from autofluorescence and motion artefacts (isosbestic 405 nm wavelength). All signals were acquired using Synapse Essentials software (Tucker Davis Technologies). Signals were sampled at 6.1 kHz (before demodulation) and 1017 Hz (after demodulation). Behavioral events (e.g., licks and sipper presentations) were time stamped by registering TTLs generated by the Med-PC system. The demodulated signals were filtered by using FFT to convert each signal from the time domain into the frequency domain, subtracting the 405 signal from the 470 signal, and then converting back into the time domain (Konanur et al., 2020). This corrected signal was expressed as a change in fluorescence, relative to total fluorescence, and used for all further analysis.

Subsequently, data were divided into discrete trials by alignment with timestamps representing the first lick in each trial and binning into 100 ms bins. Z-scores were calculated for each trial by taking the mean divided by the standard deviation of a baseline period lasting for 10 seconds preceding the first lick in each trial. Area under the curve (AUC) was calculated for the 5 seconds following the first lick before the sipper retracted and for the 5 seconds following sipper retraction. Baseline activity for each session was estimated by calculating the AUC of the epoch from the start of the session until the first trial began.

### Histology

After completion of behavioral testing and recordings, rats were deeply anaesthetized using 5% isoflurane followed by pentobarbital (50 mg/ml) before being transcardially perfused with cold 0.1 M phosphate buffered saline (PBS) followed by 4% paraformaldehyde (PFA) solution. Brains were then post-fixed overnight in ice cold 4% PFA before being transferred in 0.1 M PBS solution with 30% sucrose for at least 48 h at 4°C. Serial coronal sections (40 µm thick) were cut on a freezing microtome and stored in PBS solution containing 0.02% sodium azide. VTA-containing sections were selected to check virus spread and the position of the fiber track. Free-floating sections were transferred to 6-well plates filled with PBS. First, sections were rinsed in 0.1 M PBS (3 x 5 min) before being incubated for 1 h in blocking solution (3% goat serum, 3% donkey serum, 3% Triton in 0.1 M PBS). Next, sections were incubated overnight at room temperature with primary antibody to detect GCaMP (chicken anti-GFP, A10262, ThermoFisher Scientific; RRID: AB_2534023; 1:1000 in blocking solution). After rinses in 0.1 M PBS (3 x 5 min), sections were incubated with secondary antibody solution (goat anti-chicken IgG Alexa Fluor 488 conjugate, A-11039, ThermoFisher Scientific; RRID: AB_2534096; 1:250 in 0.1 M PBS) for 90 min at room temperature. Finally, sections were rinsed with 0.1 M PBS (3 x 5 min) and mounted in VectorShield Hard Set mounting medium and cover-slipped. Images were taken using an epifluorescence microscope (Leica DM2500) using 2.5x, 10x and 20x objectives and a R6 Retiga CCD camera (QImaging). Fiber position and virus spread were determined according to neuroanatomical landmarks (Paxinos and Watson, 1998).

### Experimental Design and Statistical Analysis

Behavioral data (lick timestamps) were extracted from data files and analyzed using custom Python scripts that measured numbers of licks for each solution and latencies from sipper extension. Position of rats in the chamber was determined using DeepLabCut (Mathis et al., 2018; Nath et al., 2019) to track body parts (nose, ears, base of tail) of rats in every frame across the preference session.

For statistical analysis of within session behavioral and neural variables, two-way mixed repeated measures ANOVA was used with Diet group as a between-subject variable (e.g. protein-restricted vs non-restricted) and Solution as a within subject variable (casein vs. maltodextrin). Choice data were analyzed by comparing diet groups using an unpaired t-test and for preference within each diet group using one-sample t-tests vs. no preference (0.5). For comparison of behavioral and neural latencies, these values were pooled for individual trials across all rats. Pearson’s correlation coefficients were calculated between latency for neural activity to peak and latency to lick (from sipper extension). Differences for each type of latency were compared between solutions using Mann-Whitney U test.

For summary data, across all sessions, two-way mixed repeated measures ANOVA was used with Diet as a between-subject variable and Session as a within-subject variable. To examine neural activity for each rat individually, AUC of casein trials was compared to AUC of maltodextrin trials using an unpaired two-tailed t-test. Resulting p-values were used to construct pie charts (threshold p<0.05).

For data from conditioning sessions, three-way mixed repeated measures ANOVA was used with Diet group as a between-subject variable (e.g. protein-restricted vs non-restricted) and Solution and Session as within subject variables (casein vs. maltodextrin; session 1 vs. session 2). For body weight, two-way mixed repeated measures ANOVA was used with Diet as a between-subject variable and Day as a within-subject variable and planned t-tests were used to compare groups on the first and last day. For food intake, unit of statistic was ‘cage’ as all rats were group housed and average food intake per rat across all days was compared with t-test.

Significant effects and interactions were followed by estimating effect sizes between subgroups. Effect sizes were determined by comparison to bootstrapped sampling distributions, which are shown in lower panels for each comparison. 5000 bootstrap samples were taken. Confidence intervals are bias corrected and accelerated and are shown on the same plots and reported in the text. Reported p-values are permutation p-values resulting from t-tests comparing 5000 reshuffles.

### Data and Software Availability

All data files are available at Figshare (doi: 10.25392/leicester.data.7636268). These experiments used a combination of software tools: Python (data extraction, analysis and plotting), and R (statistics). Estimation plots were adapted from *dabest v0.3.01* (Ho et al., 2019). All code is available at Github (https://github.com/mccutcheonlab/PPP_analysis/releases/tag/v1.0).

## Results

VTA neurons were targeted by injecting an AAV encoding the calcium sensor GCaMP6s (under control of the synapsin promoter) and a fibre optic was implanted above the injection site to record neural activity in freely moving rats (n = 14; Fig. 1A-B). Three to four weeks after surgery, a subset of rats were switched to low protein diet (5% protein from casein; PR group, n=8) while the remaining animals remained on regular chow (14% protein; NR group, n=6). Analysis of body weight data for the subsequent two weeks – before conditioning sessions started - revealed that PR and NR rats gain weight at a slightly different rate across days (Extended Data 1-1A; two-way ANOVA, Diet: F(1, 13)=0.09, p=0.767; Day: F(14, 182)=25.02, p<0.0001; Diet x Day: F(14, 182)=3.97, p<0.0001). However, the difference between diets was minimal as planned comparisons of PR and NR rats on either the first or last day did not reveal a difference in body weight between groups (Day 1: t(13)=0.72, p=0.486 and Day 14: t(13)=0.15, p=0.881). Analysis of food intake showed that PR rats exhibited a mild hyperphagia as has been previously reported (Extended Data 1-1B; mean difference in food intake between NR and PR rats: 3.77 g [95%CI 1.28, 6.88], p=0.042) (Laeger et al., 2014).

Following five days of saccharin pre-training, rats received four daily conditioning sessions in which they had access to distinctly-flavored solutions containing either casein (protein) or maltodextrin (carbohydrate; one session per day), alternated from day to day (Fig. 1B). Both groups similarly increased their consumption throughout conditioning (Extended Data 1-2; three-way ANOVA, Session: F(1,13)=22.308, p<0.0001) for both casein and maltodextrin (all Fs < 1; all Ps > 0.1). Thus, rats in both physiological states experienced the same exposure to casein and maltodextrin solutions in advance of the preference test session.

### Protein preference is associated with elevated VTA response to protein over carbohydrate

Following conditioning sessions, we then recorded VTA responses during a test session (Fig. 1B). Rats first experienced 45 trials in which only one bottle was available at a time (*forced choice trials*), similar to conditioning sessions.

Across all forced choice trials, rats exhibited similar licking behavior for casein and maltodextrin (Fig. 2A; two-way ANOVA: all Fs < 1 and all Ps > 0.1). However, PR rats did show shorter latencies to drink for casein than for maltodextrin (Fig. 2B; two-way ANOVA, Diet: F(1,13)=4.83, p=0.047; Solution: F(1,13)=9.52, p=0.009; Diet x Solution: F(1,13)=5.83, p=0.031; paired mean difference in latency between casein and maltodextrin for PR rats: - 2.48 s [95%CI -3.65, -1.03], p=0.011). In addition, PR rats on average spent more time closer to the casein sipper than the maltodextrin sipper (Extended Data 2-1; two-way ANOVA, Diet: F(1,12)=0.20, p=0.661; Solution: F(1,12)=0.50, p=0.492; Diet x Solution: F(1,12)=5.03, p=0.045; paired mean difference in distance to casein and maltodextrin sippers in NR rats: -33.2 pixels [95%CI -96.3, 59.5], p=0.375; paired mean difference in PR rats: 63.9 pixels [95%CI -11.3, 90.1], p=0.026).

**Figure 2.**
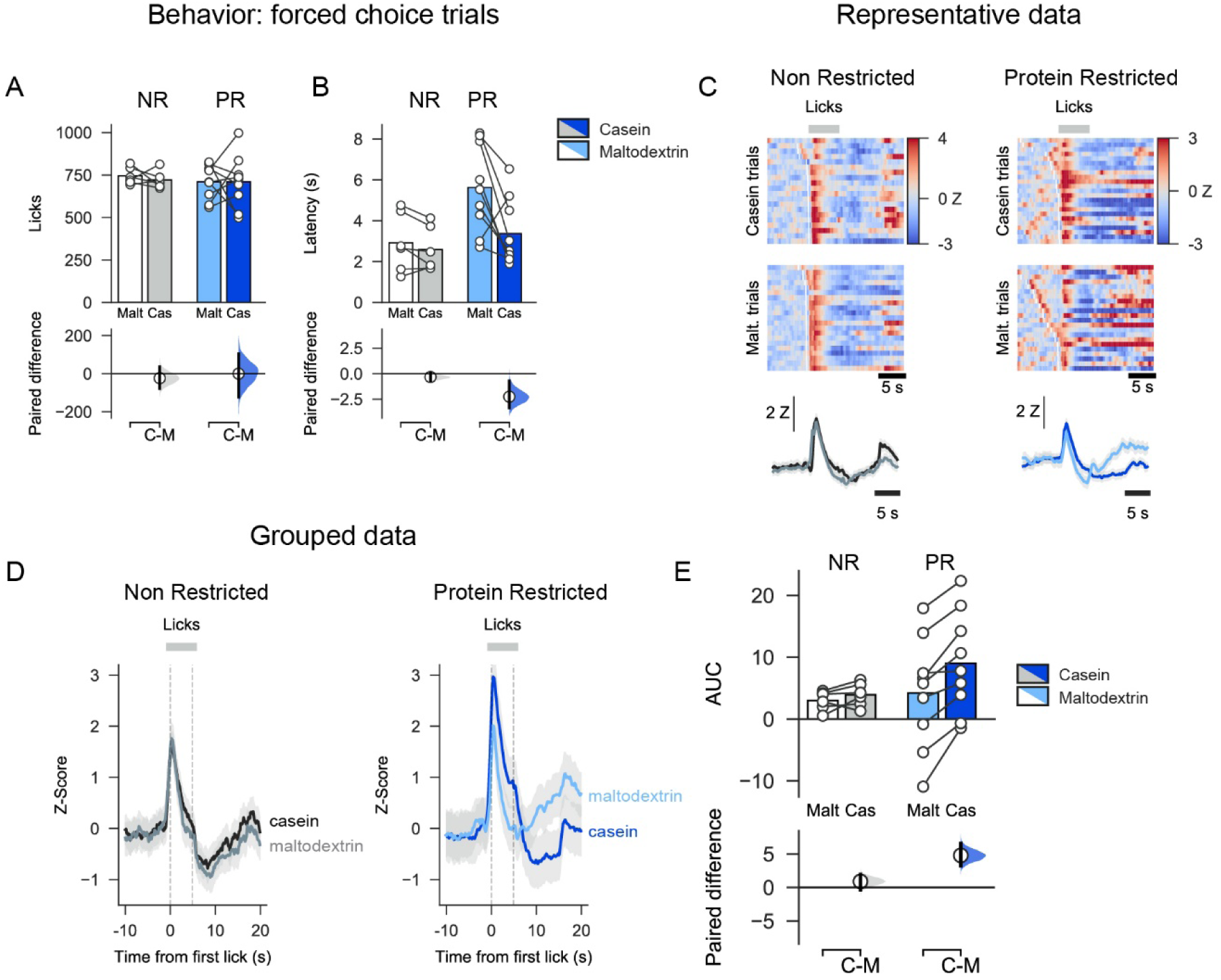
Increased neural activity in VTA of protein-restricted rats during casein consumption than maltodextrin. ***A.*** On forced choice trials, there was no difference in number of total licks for maltodextrin (Malt) vs. casein (Cas) in non-restricted (NR) or protein-restricted (PR) rats. ***B.*** Latency to drink from each sipper was influenced by diet group with PR rats showing shorter latencies on casein trials than maltodextrin trials. ***C.*** Heat maps for a single representative NR rat (left) and PR rat (right) showing normalized fluorescence changes (Z-scored) evoked by consumption of casein (top) or maltodextrin (middle) on forced choice trials. Trials are sorted by latency between sipper extension and first lick and white lines show time of sipper extension. Average fluorescence change across all trials is shown with solid line as mean and shaded area is SEM (bottom). ***D.*** Group data from forced choice casein and maltodextrin trials showing Z-score calculated from fluorescent changes aligned to first lick and averaged across all non-restricted rats (left) and protein-restricted rats (right). Solid line is mean and shaded area is SEM. ***E.*** Greater neural activation to casein consumption than maltodextrin in PR rats but not NR rats as shown by area under curve (AUC, 0-5 seconds following first lick). In A, B, and E, upper panels show mean as bars and data from individual rats as circles while lower panels show mean difference as a bootstrap sampling distribution with mean differences depicted as dots and 95% confidence intervals indicated by the ends of the vertical error bars.

Photometry recordings of VTA neurons during consumption of each solution (Fig. 2C-E) showed that casein and maltodextrin consumption evoked similar VTA responses in NR rats (paired mean difference in AUC between casein and maltodextrin in NR rats: 0.80 [95%CI - 0.46, 2.17], p=0.354). In contrast, although PR rats licked similarly for both solutions (Fig. 2A), casein consumption is associated with a higher VTA response than for maltodextrin (Fig. 2E; two-way ANOVA, Diet: F(1,13)=0.60, p=0.454; Solution: F(1,13)=20.73, p=0.0005; Diet x Solution: F(1,13)=10.39, p=0.007; paired mean difference in AUC between casein and maltodextrin in PR rats: 4.66 [95%CI 3.27, 6.41], p=0.0026). No differences were found in neural activity in the 5 s epoch following termination of licking (Extended Data 2-2; two-way ANOVA, Diet: F(1,13)=0.96, p=0.346; Solution: F(1,13)=1.80, p=0.203; Diet x Solution: F(1,13)=0.05, p=0.824). Moreover, differences in VTA responses were not attributable to differences in baseline activity between the two diet conditions (Extended Data 2-3; unpaired t-test: t(13)=0.30, p=0.769).

We examined whether there were differences in how long the photometry signal took to peak during each trial and whether this was correlated with the latency to lick (Extended Data 2-4). We found that in NR rats there was no difference between casein and maltodextrin trials in latency to peak calcium response (from sipper extension; Mann-Whitney U: p=0.743). Moreover, on a trial-to-trial basis the latency to peak showed a moderate but significant correlation with latency to lick for both casein trials (Pearson correlation coefficient: r=0.27, p=0.0014) and maltodextrin trials (r=0.20, p=0.020).

In contrast, for PR rats, the latency for the photometry signal to peak did differ between casein and maltodextrin trials (Mann-Whitney U: p<0.001) and, furthermore, there was a highly significant correlation with latency to lick on casein trials (r=0.45, p<0.0001), but no correlation for maltodextrin trials (r=0.11, p=0.148). These findings for PR rats are likely due to the neural activation at time of licking on maltodextrin trials being greatly reduced for this group of rats.

Following these forced choice trials, rats were presented with twenty trials in which both bottles were available at the same time (*free choice trials*) to confirm the existence of protein preference in the PR group (Murphy et al., 2018; Naneix et al., 2019). In free choice trials, PR rats significantly licked more for casein than for maltodextrin (Fig. 3A; two-way ANOVA, Diet: F(1,13)=5.12, p=0.041; Solution: F(1,13)=1.75, p=0.208; Diet x Solution: F(1,13)=14.96, p=0.002; mean paired difference in licks between casein and maltodextrin for PR rats: 442.22 [95%CI 127.33, 587.22], p=0.006), whereas NR rats did not (mean paired difference in licks between casein and maltodextrin for NR rats: -216.67 [95%CI -464.67, -16.16], p=0.121). Consistent with this result, PR and NR rats exhibited differential casein preference, as calculated by the number of times they chose casein during the free choice trials (mean difference in choice preference between NR and PR rats: difference between groups: 0.49 [95%CI 0.23, 0.66], p=0.004). As such, NR rats showed no preference for one solution over the other (preference for NR rats: 0.37 [95%CI 0.23, 0.52], p=0.121 vs. 50%) but PR rats displayed a strong preference for casein (Fig. 3B; preference for PR rats: 0.85 [95%CI 0.58, 0.95], p=0.0064 vs. 50%).

**Figure 3.**
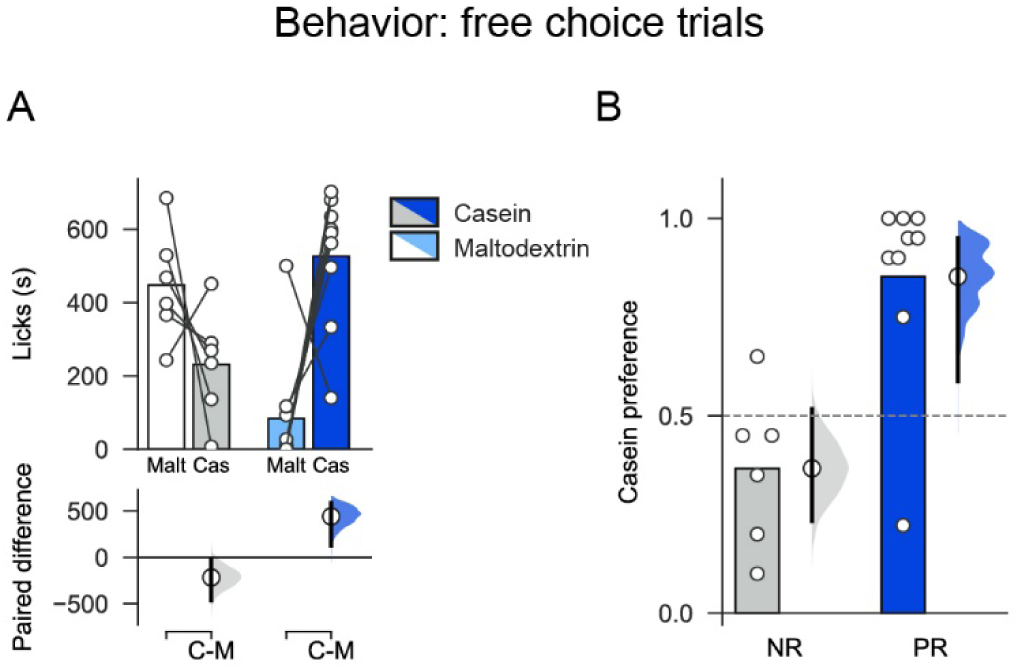
Protein-restricted rats show a strong preference for protein over carbohydrate that is not seen in control rats. ***A,*** On free choice trials, protein-restricted (PR) rats licked more than casein than maltodextrin but there was no difference in licking between the solutions in non-restricted (NR) rats. ***B,*** When number of choices for each solution were considered, PR rats showed a strong preference for casein relative to maltodextrin. Bars show mean and circles are data from individual rats. Bootstrapped sampling distributions are used to show mean paired difference in lower panel of A and difference vs. 0.5 to the right of bars in B. Means of distributions are shown as dots and 95% confidence intervals indicated by the ends of the vertical error bars.

### Preference towards Protein Develops with Minimal Experience in a Newly Protein Restricted State

Next, we were interested in what would happen to behavior and neural activity when rats’ protein needs changed. First, we investigated what happened when rats from the control group were switched to the protein-restricted diet (hereafter, NR → PR rats). Importantly, we re-tested rats at two time points: one week after diets were switched but before any intervening experience of the casein and maltodextrin solutions (Fig. 4A; Pref. Test 2) and one week after this, once rats had experienced an extra block of conditioning sessions (Fig. 4G; Pref. Test 3).

**Figure 4.**
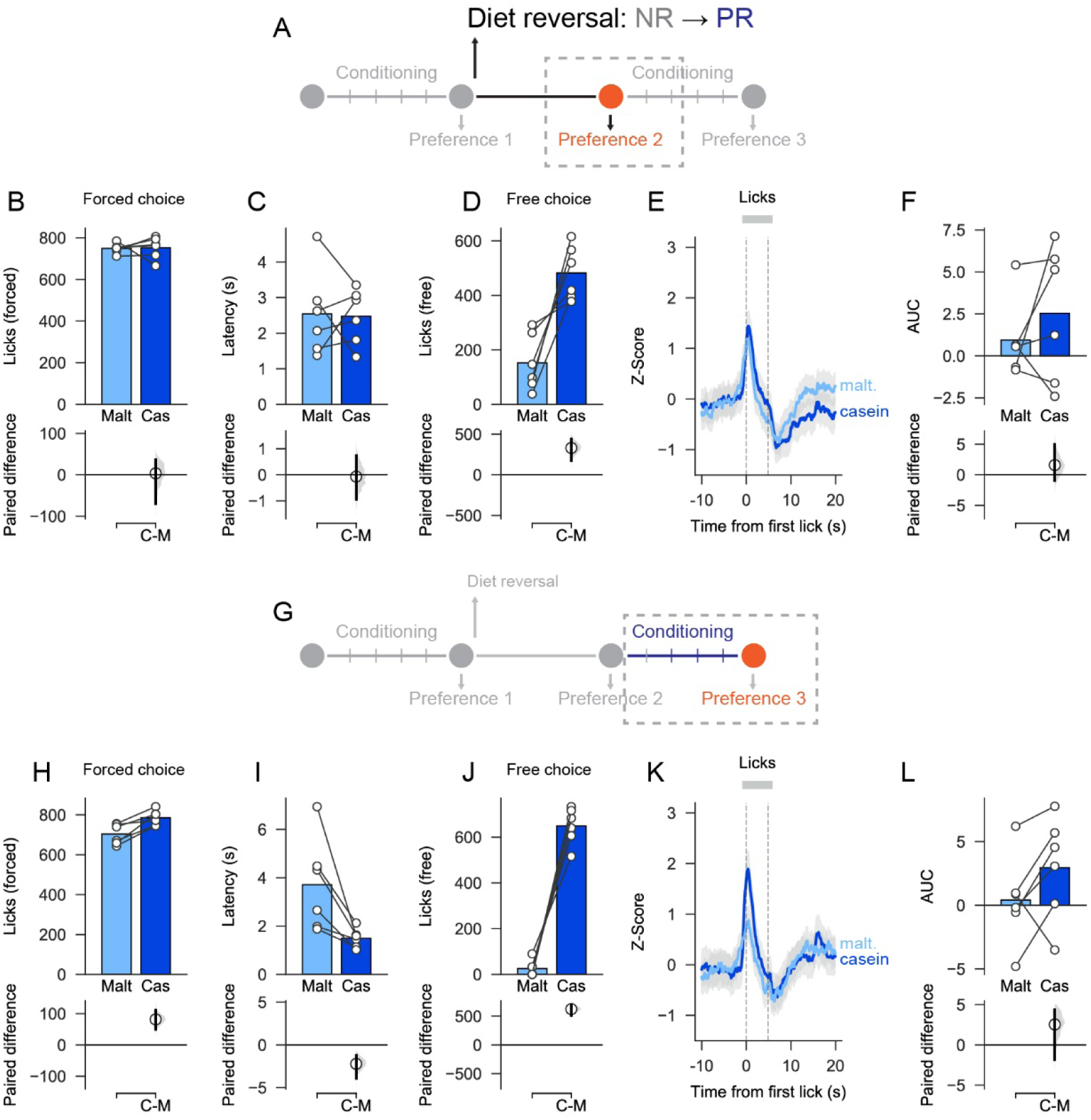
Changing from control diet to low protein diet leads to changes in behavior toward nutrients. ***A,*** Schematic showing experimental timeline for Preference Test 2 (before additional conditioning sessions). ***B-C,*** On forced choice trials, there was no difference in licks for casein and maltodextrin or in latency to drink from each sipper. ***D,*** On free choice trials, rats licked more for casein than maltodextrin. ***E-F,*** As a group, VTA neural activity was similar between casein and maltodextrin trials but there was a large amount of variability. ***G,*** Schematic showing experimental timeline for Preference Test 3 (after additional conditioning sessions). ***H-I,*** On forced choice trials, there was a small increase in licks for casein relative to maltodextrin and latency to drink was shorter on casein trials than maltodextrin trials. ***J,*** On free choice trials, rats licked more for casein than maltodextrin. ***K-L,*** VTA neural activity was not different between casein and maltodextrin trials although, as with the previous test, there was a high degree of variability. Upper panels show mean as bars and data from individual rats as circles while lower panels show mean difference as a bootstrap sampling distribution with mean differences depicted as dots and 95% confidence intervals indicated by the ends of the vertical error bars.

As reported in Pref Test 1 (see above), animals licked similarly for casein and maltodextrin during forced choice trials in Pref Test 2 (Fig. 4B; mean paired difference in licks between casein and maltodextrin: 3.5 [95%CI -69.5, 36.0], p=0.817) but slightly increased the licking for casein in Pref Test 3 (Fig. 4H; mean paired difference: 81.50 [95%CI 50.00, 111.17], p<0.001). Similarly, analysis of latencies indicated no difference during Pref Test 2 (Fig. 4C; mean paired difference in latency between casein and maltodextrin: -0.07 s [95%CI -0.94, 0.73] p=0.974), but showed shorter latencies to drink from the casein sipper during Pref Test 3 (Fig. 4I; mean paired difference: -2.22 s [95%CI -3.91, -1.23], p=0.030).

On free choice trials NR → PR rats licked more for casein than maltodextrin during both Pref Test 2 (Fig. 4D; mean paired difference in licks between casein and maltodextrin: 330.00 [95%CI 176.33, 440.17], p<0.001) and Pref Test 3 (Fig. 4J; mean paired difference: 623.17 [95%CI 511.17, 689.83], p<0.001). As expected, this pattern resulted in strong casein preference over maltodextrin on Pref Test 2 (preference: 0.71 [95%CI 0.60, 0.83], p=0.030 vs. 50%) and Pref Test 3 (preference: 0.95 [95%CI 0.83, 0.98], p=0.030 vs. 50%).

The casein preference reported in Pref Test 2 and Pref Test 3 in NR → PR rats strongly contrasts with behavior during the first preference test (Fig. 3). Interestingly, photometry recordings during forced choice trials did not show any difference in VTA responses to casein and maltodextrin in either Pref Test 2 (Fig. 4E-F; mean paired difference in AUC between casein and maltodextrin: 1.59 [95%CI -0.92, 4.95] p=0.381) or Pref Test 3 (Fig. 4K-L; mean paired difference: 2.53 [95%CI -1.84, 4.37], p=0.097).

In summary, NR → PR rats developed a rapid behavioral preference to protein over carbohydrate that was observed even before they had gained extensive experience with each solution. Activity in VTA, however, was slower to change to reflect the rats new physiological state and behavior.

### Protein Preference and Differences in Associated VTA Activity Disappear After Experience with Nutrient Solutions In Protein Replete State

We also investigated the effect of protein repletion on casein preference and VTA responses using a similar diet switch design in rats that were initially protein restricted were changed to non-restricted diet (hereafter, PR → NR rats). Again, rats were tested one week following the diet switch but before being given additional experience with solutions (Pref Test 2; Fig. 5A) and then, again, after a block of conditioning sessions (Pref Test 3; Fig. 5G).

**Figure 5.**
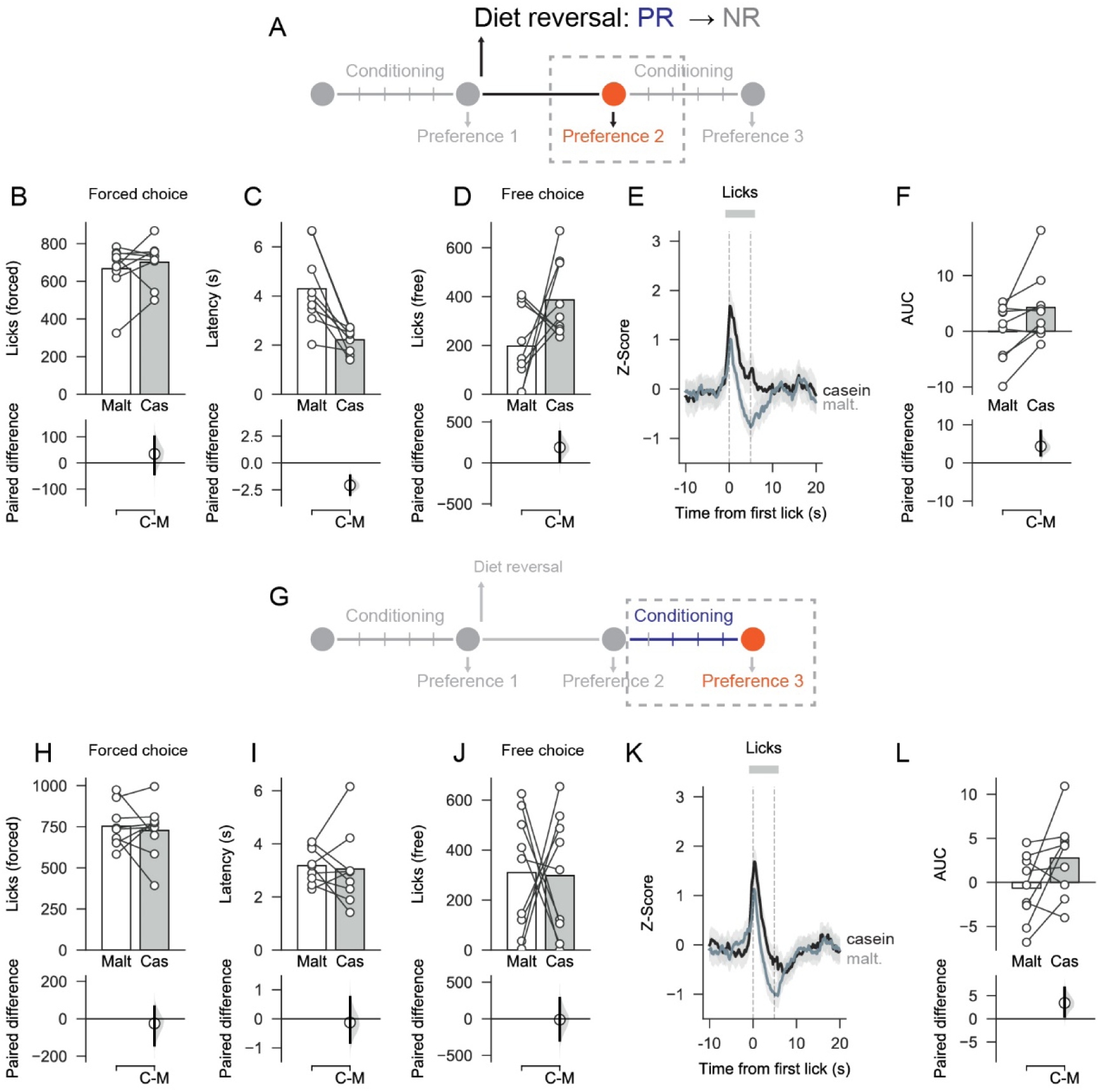
Changing from low protein diet to control diet leads to changes in behavior toward nutrients. ***A,*** Schematic showing experimental timeline for Preference Test 2 (before additional conditioning sessions). ***B-C,*** On forced choice trials, there was no difference in licks for casein and maltodextrin but latency to drink was shorter on casein trials than on maltodextrin trials. ***D,*** On free choice trials, number of licks was similar for casein and maltodextrin although rats chose the casein sipper more than the maltodextrin (see Results). ***E-F,*** VTA neural activity was elevated on casein trials vs. maltodextrin trials. ***G,*** Schematic showing experimental timeline for Preference Test 3 (after additional conditioning sessions). ***H-I,*** On forced choice trials, the number of licks and latencies were similar for casein and maltodextrin trials. ***J,*** On free choice trials, number of licks was similar for casein and maltodextrin. ***K-L,*** VTA neural activity was no longer different between casein and maltodextrin trials. Upper panels show mean as bars and data from individual rats as circles while lower panels show mean difference as a bootstrap sampling distribution with mean differences depicted as dots and 95% confidence intervals indicated by the ends of the vertical error bars.

During forced choice trials there was no difference in the number of licks for casein and maltodextrin in Pref Test 2 (Fig. 5B; mean paired difference in licks between casein and maltodextrin: 34.67 [95%CI -42.44, 100.44], p=0.386) or Pref Test 3 (Fig. 5H; mean paired difference: -25.00 [95%CI -141.78, 64.33], p=0.682). The latency to drink from the casein sipper was still shorter than the latency for maltodextrin in Pref Test 2 (Fig. 5C; mean paired difference in latency between casein and maltodextrin: -2.11 s [95%CI -2.95, -1.22], p=0.003) but this difference disappeared in Pref Test 3 after additional conditioning sessions (Fig. 5I mean paired difference: -0.24 s [95%CI -0.91, 0.65], p=0.561).

On free choice trials there was now no significant difference in the number of licks between casein and maltodextrin during Pref Test 2 (Fig. 5D; mean paired difference in licks between casein and maltodextrin: 189.22 [95%CI 19.89, 380.44], p=0.099) although when number of choices was considered, as a group, PR → NR rats still showed a moderate preference for casein over maltodextrin (preference: 0.68 [95%CI 0.57, 0.79], p=0.020 vs. 50%). In Pref Test 3 after additional conditioning sessions, casein preference was completely abolished for both licking (Fig. 5J; mean paired difference in licks between casein and maltodextrin: -11.78 [95%CI -294.11, 279.67], p=0.922) and choices (preference: 0.48 [95%CI 0.26, 0.68], p=0.889 vs. 50%).

When VTA neural activity was analyzed during forced choice trials we found that there was still greater VTA activation on casein trials than maltodextrin trials during Pref Test 2 although the effect size was more variable than on the first preference test (Fig. 5E-F; mean paired difference in AUC between casein and maltodextrin 3.86 [95%CI 1.54, 8.17], p=0.028). Consistent with the abolition of casein preference reported during Pref Test 3, analysis of VTA neural activity also now showed no reliable difference between casein and maltodextrin in forced choice trials although there was a high degree of variability (Fig. 5K-L; mean paired difference in AUC between casein and maltodextrin 3.24 [95%CI 0.47, 6.37], p=0.091). Thus, the protein preference and associated VTA responses that developed when rats were protein-restricted was markedly reduced once rats had gained additional experience with the nutrient solutions in the new protein replete state.

### Behavior and VTA Activity Become Uncoupled after Diet Switch

To compare across all sessions for each group of rats, we examined how protein preference changed from preference test 1 to test 3. After the switch from non-restricted to protein-restricted state (NR → PR rats), there was a clear shift in behavior across the three sessions as shown by a main effect of Session (Fig. 6A; one-way repeated measures ANOVA: F(2,10)=27.01, p<0.0001). Further comparisons showed that after diet switch NR → PR rats’ behavior differed both before additional conditioning sessions (mean paired difference in preference between Pref. Test 2 and Test 1: 0.34 [95%CI 0.16, 0.52], p=0.007) and after (mean paired difference between Pref. Test 3 and Test 1; 0.58 [95%CI 0.43, 0.73], p=0.001). However, consistent with our earlier analysis, VTA responses to casein and maltodextrin did not significantly change between the three preference tests (Fig. 6B; two-way repeated ANOVA: Session (F(2,10)=0.72, p=0.508; Solution (F(1,5)=2.07, p=0.21); Session x Solution (F(2,10)=3.02, p=0.094)).

**Figure 6.**
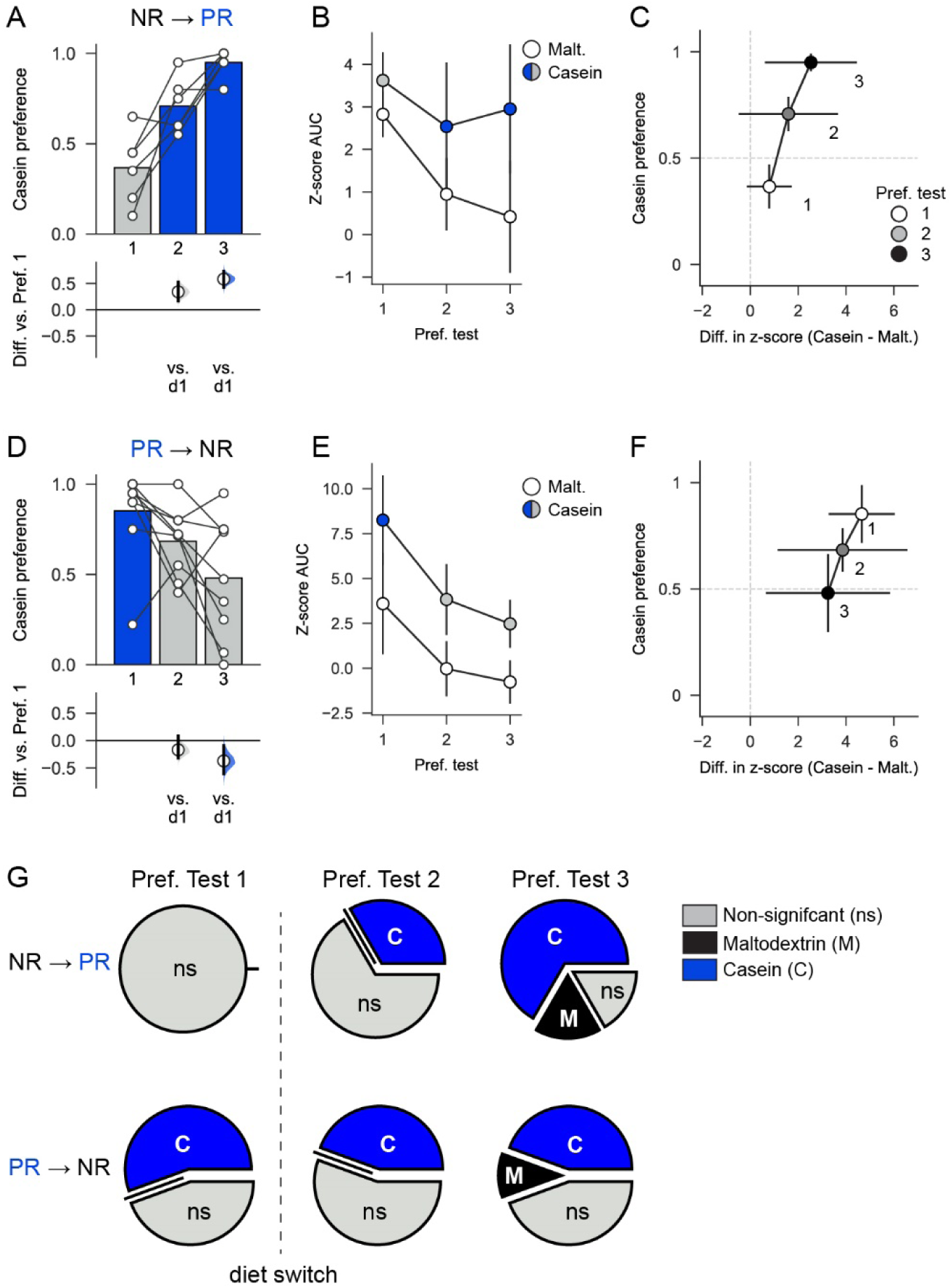
Behavior and VTA activity become uncoupled after diet switch. ***A,*** In NR → PR rats, preference for protein increases after diet switch in both preference test 2, without additional conditioning, and in preference test 3. Bars are mean and circles show data from individual rats with mean differences of bootstrapped sampling distributions shown in lower panel vs. preference test 1. ***B,*** Neural activity in VTA on casein and maltodextrin trials is not affected by diet switch. ***C,*** Behavioral preference for casein vs. maltodextrin (y-axis) plotted as a function of difference in neural activation (z-score AUC) associated with consumption of each solution (x-axis) in NR → PR rats. Circles connected by black solid lines show mean ± SEM. ***D,*** In PR → NR rats, behavior changes after diet switch but requires additional conditioning sessions for protein preference to shift, relative to preference test 1. Plotting conventions as in A. ***E,*** Neural activity in VTA is consistently elevated on casein trials, relative to maltodextrin trials. ***F,*** Preference vs. difference in neural activation for PR → NR rats with plotting conventions as in C. ***G,*** Neural activity evoked by consumption of each solution changes as a function of diet state. Pie charts show the proportion of rats showing significantly greater activation to maltodextrin (black) or casein (blue) with non-significant shown in grey. Upper panel, For non-restricted (NR) rats, there is no difference in neural activity between casein and maltodextrin on the first preference test whereas after diet switch a progressively greater number of rats show a preference for casein. Lower panel, For protein-restricted (PR) rats, a majority show greater activation to casein than to maltodextrin and even after switching to control diet 4 out of 9 rats continue to show greater VTA activation to casein than to maltodextrin.

In contrast, protein repletion (PR → NR rats) induced a gradual decrease in casein preference across the three tests (Fig. 6D; one-way repeated ANOVA: F(2,16)=5.99, p=0.011). Between sessions comparisons showed that casein preference in second test session, when rats had not received additional conditioning, was no different to the first test session (mean paired difference in preference between Pref. Test 2 and Test 1: -0.17 [95%CI -0.31, 0.09], p=0.119). However, by the third test session there was a significant decrease in casein preference compared to the first session (mean paired difference in preference between Pref. Test 3 and Test 1: -0.37 [95%CI -0.60, -0.09], p=0.018). This shift in casein preference is associated with a trend towards a decrease in VTA responses to both casein and maltodextrin through the three sessions (Fig. 6E; two-way repeated ANOVA: Session F(2,16)=4.57, p=0.06; Session x Solution F(2,16)=0.42, p=0.666). However, VTA responses to casein remained higher than responses to maltodextrin (Solution: F(1,8)=14.25, p=0.005).

The relationship between casein preference and neural activation to each solution is summarized in Fig. 6C and 6F. Performing a simple linear regression between behavior (casein preference) and photometry (difference in z-score between casein and maltodextrin trials) yielded weak-to-moderate correlations for each group with this relationship being significant for PR → NR rats (Pearson’s correlation, r=0.41, p=0.034; Fig. 6C) but not for NR → PR rats (r=0.23, p=0.350; Fig. 6F).

Next, we performed multivariate linear regression on these data including test day as a predictor and found higher beta coefficients associated with behavior than with photometry supporting our finding that protein preference changed more readily across the dietary manipulations than did neural activity (beta coefficients for behavior: 2.51 and -1.55 for NR → PR rats and PR → NR rats, respectively; beta coefficients for photometry: 0.02 and 0.02 for NR → PR rats and PR → NR rats, respectively). In addition, beta coefficients for behavior were oppositely signed in each diet group reflecting the bidirectional change in behavior.

Finally, to check whether behavior and photometry measurements were more closely related to state of protein deprivation we re-coded data based on each animal’s current dietary state and re-ran the regression. Once again we found that higher beta values were associated with behavior than with photometry (behavior: 1.32 and 0.76 for NR → PR rats and PR → NR rats, respectively; photometry: -0.01 and 0.01 for NR → PR rats and PR → NR rats, respectively).

These analyses and visual inspection of the data suggested that changes in VTA responses after diet switch may have been obscured by inter-individual variability in responses. To explore this further, we chose to look at differences in VTA activity on a rat-by-rat basis. By comparing activity on individual trials – rather than the mean of these trials – we calculated for each rat whether there was significantly greater activation to casein or to maltodextrin (Fig. 6G). For NR → PR rats, no rats showed a significantly greater activation to either nutrient on Pref. Test 1. However, after switching to the protein-restricted diet a progressively greater proportion showed significantly greater activation to casein (33% for Pref. Test 2, 66% for Pref. Test 3). For PR → NR rats, results were strikingly different. On Pref. Test 1 a majority of rats (56%) showed significantly greater activation on casein trials than on maltodextrin trials. After switching to control diet this changed little, with a large proportion continuing to show greater activation on casein than on maltodextrin trials (44% on both Pref. Test 2 and Pref. Test 3).

In summary, protein preference behavior changed strongly and rapidly in a bidirectional manner in both groups of rats while shifts in VTA neural activity were not as apparent especially in PR → NR rats.

## Discussion

Animals prioritize protein intake over the intake of other macronutrients (Morrison and Laeger, 2015). However, the neural mechanisms underpinning this behavioral process are not well understood. Here, for the first time, we show that protein restriction changes neural activity in the VTA during the consumption of protein or carbohydrate to reflect the initial protein preference. Furthermore, we also demonstrate that protein preference is dependent on current physiological state and can be induced or abolished after according to protein needs. Interestingly, VTA nutrient-related responses are highly dependent on the animal’s prior experience in protein restricted or non-restricted state, appearing slower than behavior to adapt to new physiological status.

### Protein appetite is associated with increased VTA activity

Consistent with our earlier studies (Murphy et al., 2018; Naneix, Peters, McCutcheon, 2019), protein-restricted rats developed a strong preference for protein-containing solution over carbohydrate-containing solution. Protein preference did not coincide with a general aversion to other carbohydrate as rats consumed similar amounts of both casein and maltodextrin during conditioning and forced choice trials. This differs from responses seen to diets lacking single amino acids that can lead to development of conditioned taste aversion for foods with imbalanced amino acid content (Maurin et al., 2005; Gietzen and Aja, 2012).

VTA neurons play a complex role in the control of food-related behaviors (Berridge, 2007; Bromberg-Martin et al., 2010; Brown et al., 2012; Zessen et al., 2012; Root et al., 2020). Previous studies show that dopamine signaling originating in the VTA is involved in establishing carbohydrate-based flavor preferences (Sclafani et al., 2011; de Araujo et al., 2012; McCutcheon, 2015; Hsu et al., 2018). Here, we show for the first time that protein appetite involves VTA circuits and that VTA activation is modulated by both the macronutrient content of the food and the rats’ protein status during the initial preference test (Fig. 2). Specifically, VTA responses are greater during consumption of protein (casein) compared to carbohydrate (maltodextrin) selectively in protein-restricted rats. These differences in VTA activity are observed during forced choice trials, in which only one solution is available, but this difference in activity reflects future food preference in the subsequent free choice trials. Importantly, this difference is not the result of different behavioral activation as rats exhibited similar levels of licking. Differences in VTA responses to the consumption of each nutrient may reflect reward value and be used to guide food preferences (Berridge, 2007; Roitman et al., 2008; Bromberg-Martin et al., 2010; McCutcheon et al., 2012b; Salamone and Correa, 2012). In addition, protein-restricted rats exhibited a shorter latency for casein consumption (Fig. 2) suggesting an increase in incentive properties of this solution (Barbano and Cador, 2005). We previously reported that protein appetite was associated with increased casein palatability (Murphy et al., 2018; Naneix et al., 2019).

Using *ex vivo* voltammetry recordings, we recently showed that protein restriction increased evoked dopamine release in the nucleus accumbens, but not dorsal striatum (Naneix et al., 2020). Similar changes have been reported with other nutrients (McCutcheon, 2015) and hunger states (Heffner et al., 1980), which may be used to reinforce and guide food-seeking behaviors toward the most relevant source of food. While firing of dopamine neurons does not always reflect terminal release (Sulzer et al., 2016; Mohebi et al., 2019), this result is consistent with our present *in vivo* observation in protein-restricted rats. There is now a need to characterize if these increased VTA responses also translate into increased dopamine release *in vivo*, precisely where this release occurs in the forebrain, and how dopamine cell bodies or terminals may be able to detect dietary amino acids (Karnani et al., 2011).

### VTA responses do not follow changes in initial protein preferences

Changes in protein status after an initial nutrient preference resulted in different behavioral adaptations depending on the direction of diet shift. Rats experiencing a new protein deficiency (NR→PR; Fig. 4) rapidly shifted their preference toward casein even without additional conditioning, suggesting that protein appetite can manifest independently of prior experience with protein-containing food in a restricted state. Previous studies have demonstrated that an immediate specific appetite exists for another essential nutrient, sodium (Krause and Sakai, 2007). As such, sodium depletion induces immediate and unlearned alterations in how sodium is perceived and how animals respond to stimuli previously associated with sodium (Robinson and Berridge, 2013). However, sodium appetite is rapidly terminated once sodium levels are restored (Krause and Sakai, 2007). Such fine regulation was not observed with protein intake (PR→NR; Fig. 5) as casein preference only decreased in newly protein-replete rats after experiencing additional conditioning sessions.

VTA responses to both casein and maltodextrin became more complex and did not immediately follow changes in protein preference. Newly protein-restricted rats (NR→PR; Fig. 4) exhibited delayed changes in VTA responses to casein and maltodextrin consumption, despite increased preference for protein. Previous studies have shown that unconditioned VTA dopamine responses to food or specific nutrients (Cone et al., 2014, 2016) update immediately, independently of prior experience of the physiological state (*e.g.* sodium depletion, hunger). In contrast, dopamine responses to food- or nutrient-predictive cues require multiple associations under physiological conditions in which the food is rewarding (Bassareo and Di Chiara, 1997; Day et al., 2007; Cone et al., 2016). Thus, our results suggest that VTA activity may track the value of the flavor paired with protein rather than the protein content itself (Sclafani et al., 2011; McCutcheon, 2015).

Protein repletion (PR→NR; Fig. 5) had a delayed impact on VTA activity, as rats continued to show elevated VTA responses to casein despite a progressive decrease of their protein preference. These results contrast starkly with those from studies of sodium appetite where VTA dopamine responses to conditioned cues are flexibly expressed in a state-dependent manner once learned (Cone et al., 2016). Instead, elevated VTA responses to casein even after the initial behavioral preference was reversed, suggests a long-lasting neurobiological impact of protein restriction that may require extended time and prolonged learning to be reversed.

### Methodological considerations

In this study we used a targeting strategy that was not selective for dopamine neurons. As such, it is likely that some of the photometry signal resulted from activity in non-dopamine populations of VTA neurons including local GABA interneurons and projecting GABA or glutamate neurons (Dobi et al., 2010; Morales and Margolis, 2017) although, by number, dopamine neurons represent the largest proportion of VTA neurons (Nair-Roberts et al., 2008). In addition, the increases in neural activity evoked by behavioral events are qualitatively similar to those others have observed when recording only dopamine neurons (e.g. with TH::Cre rats; Parker et al., 2016) or when recording dopamine release using voltammetry (Phillips et al., 2003). As other VTA neuronal populations are involved in different aspects of food-related behaviors (Brown et al., 2012; Zessen et al., 2012; Morales and Margolis, 2017; Root et al., 2020), future cell-specific targeting will be required to tease apart responses from these neuronal subtypes.

This study used only male rats, consistent with our previous study (Murphy et al., 2018). Protein (and other macronutrient) requirements differ in male and female rats at adulthood and through development (Leibowitz et al., 1991) and, in addition, total food intake changes across the estrus cycle with resulting effects on the proportion of protein intake (Wurtman and Baum, 1980). Moreover, physiological state influences the activity of VTA neurons in a sex-dependent manner (Godfrey and Borgland, 2020). Thus, a better understanding of brain mechanisms underlying protein appetite warrants further investigation in both males and females.

### Conclusions

A key remaining question is how VTA midbrain circuits detect the nutrient content of food and integrate this with physiological state to regulate protein homeostasis. Previous work suggests that the VTA must receive taste information (Hajnal et al., 2004; Roitman et al., 2008; McCutcheon et al., 2012b). Protein can be detected via umami receptors expressed on taste buds (Chaudhari et al., 2009; Liman et al., 2014) but the link between protein sensing by the tongue and VTA neuronal populations remains to be explored. VTA circuits are also sensitive to the caloric content of food (de Araujo et al., 2008; Domingos et al., 2011; Beeler et al., 2012; Ferreira et al., 2012; McCutcheon et al., 2012a) and this information is relayed to forebrain regions controlling food-seeking behaviors (Tellez et al., 2016). Whether VTA neurons are sensitive to protein or amino acids directly is not known but individual amino acid levels can be detected by hypothalamic, cortical, and hindbrain regions connected to the VTA (Karnani et al., 2011; Anthony and Gietzen, 2013; Heeley and Blouet, 2016; Tsang et al., 2020). Furthermore, recent work showed that fibroblast growth factor 21 (FGF21), a hepatic hormone, is released in response to reduction in dietary protein (Laeger et al., 2014) and its central action is necessary for development of protein preference in mice (Hill et al., 2019).

Given the potential effects of inadequate protein diet *in utero* or after birth on neurodevelopmental disorders (Grissom and Reyes, 2013; Gould et al., 2018) and obesity (Simpson and Raubenheimer, 2005), our results highlight neurobiological substrates that may underlie protein appetite in normal and pathological conditions

## Acknowledgements

The authors acknowledge the help and support from the staff of the Division of Biomedical Services, Preclinical Research Facility, University of Leicester, for technical support and the care of experimental animals. The authors would like to thank Vaibhav Konanur for developing the analytical method used to correct fluorescence traces, Leon Lagnado for kindly loaning equipment used in initial photometry experiments, and Andrew MacAskill for useful discussions regarding analysis. This work was funded by the Biotechnology and Biological Sciences Research Council [grant #BB/M007391/1 to J.E.M.], the European Commission [grant #GA 631404 to J.E.M.], The Leverhulme Trust [grant #RPG-2017-417 to J.E.M. and J.A-S.], and Tromsø Research Foundation [grant #19-SG-JMcC to J. E. M.).

## Author Contributions

Conceptualization, J.E.M.; Formal Analysis, G.C., F.N. and J.E.M.; Investigation, G.C., F.N., K.Z.P., E.M.S.S. and J.E.M.; Writing – Original Draft, F.N. and J.E.M.; Writing – Review & Editing, G.C., F.N., K.Z.P., J.A-S., E.M.S.S. and J.E.M.; Funding Acquisition, J.E.M., J.A-S.

## Extended Data

**Extended Data 1-1.**
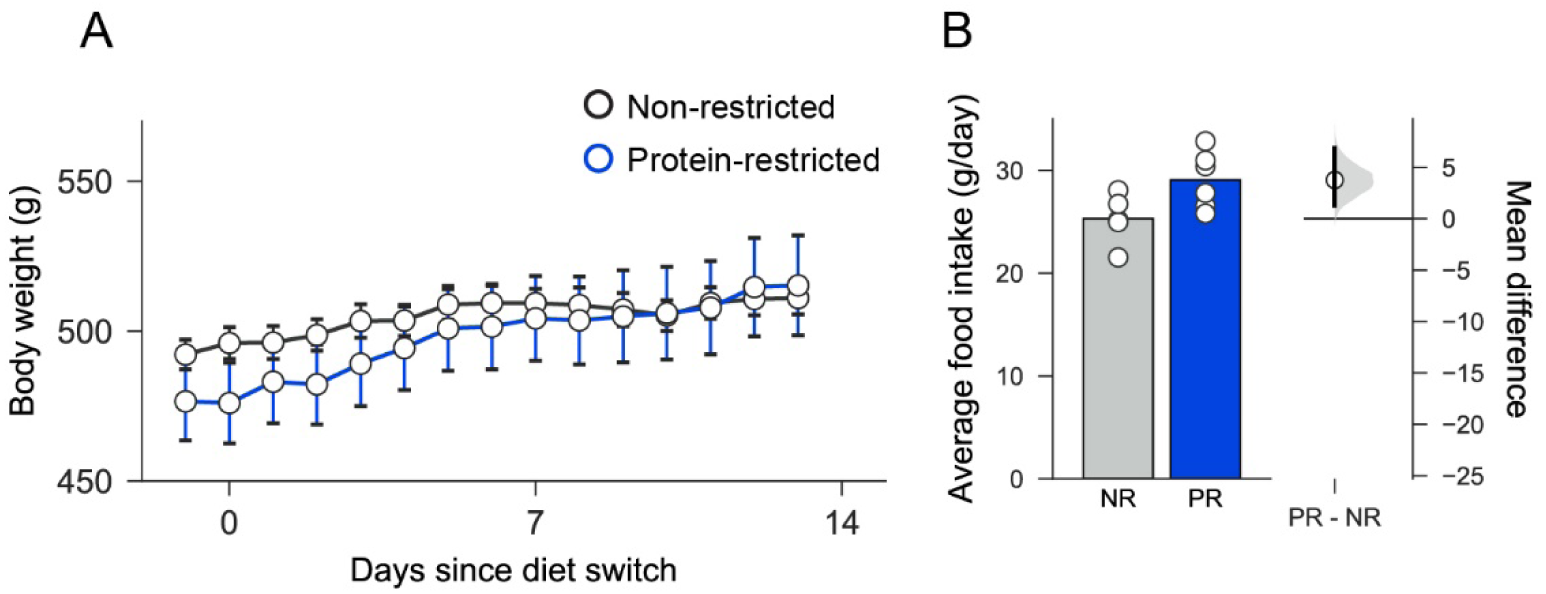
Body weight and food intake data. ***A***, Similar changes in body weight increase were seen in protein-restricted (PR) and non-restricted (NR) control rats. Circles show mean for each day and error bars are SEM. ***B***, Mild increase in food intake was seen in PR rats relative to NR rats. Left panel, bars are mean and circles are individual data points (cages). Right panel, mean difference as a bootstrap sampling distribution with mean difference depicted as dot and 95% confidence intervals indicated by the ends of the vertical error bars.

**Extended Data 1-2.**
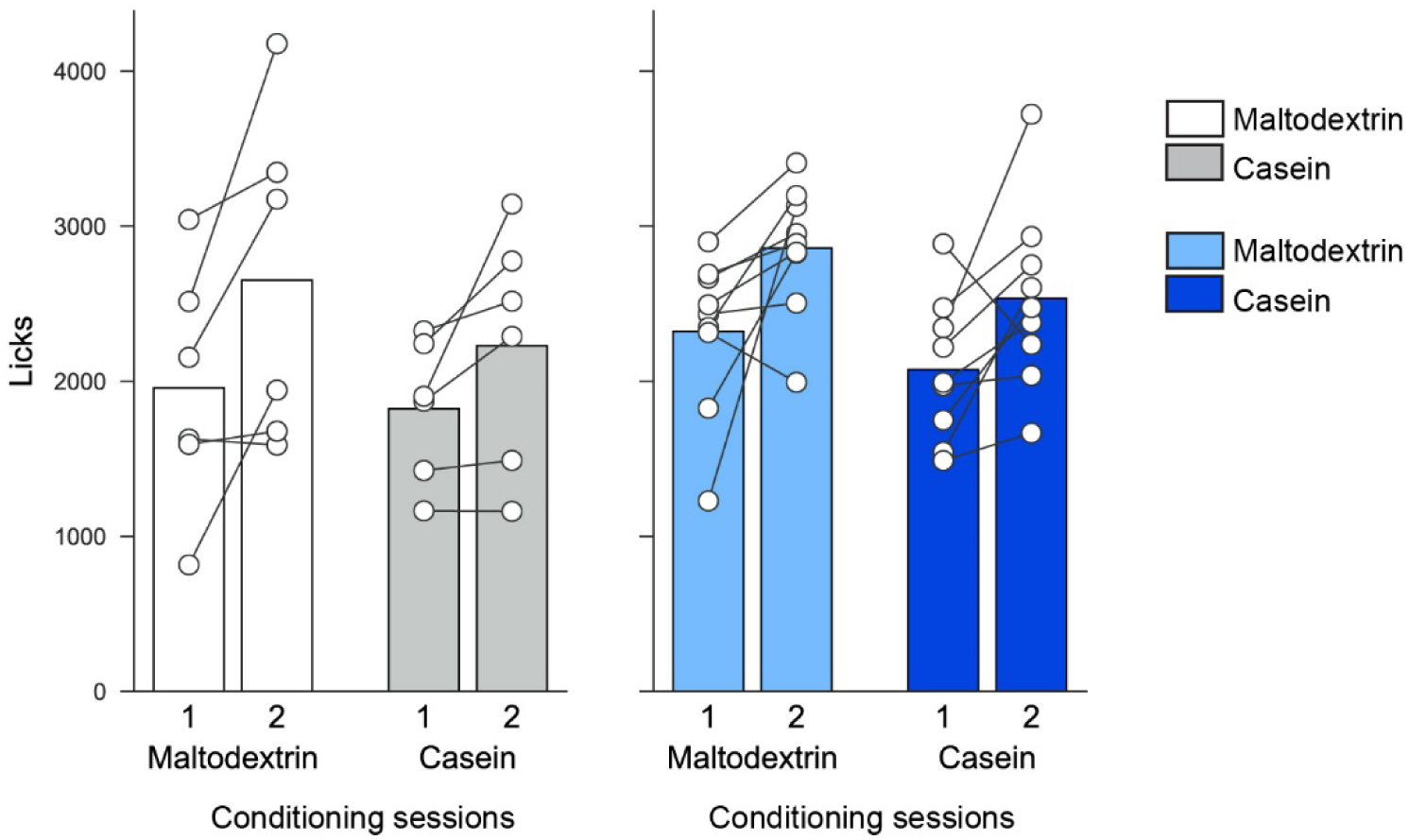
Data from conditioning sessions show that for both solutions more was consumed on the second conditioning day than on the first day but there were no differences between diet groups or solutions. Bars are mean and circles are individual data points (rats).

**Extended Data 2-1.**
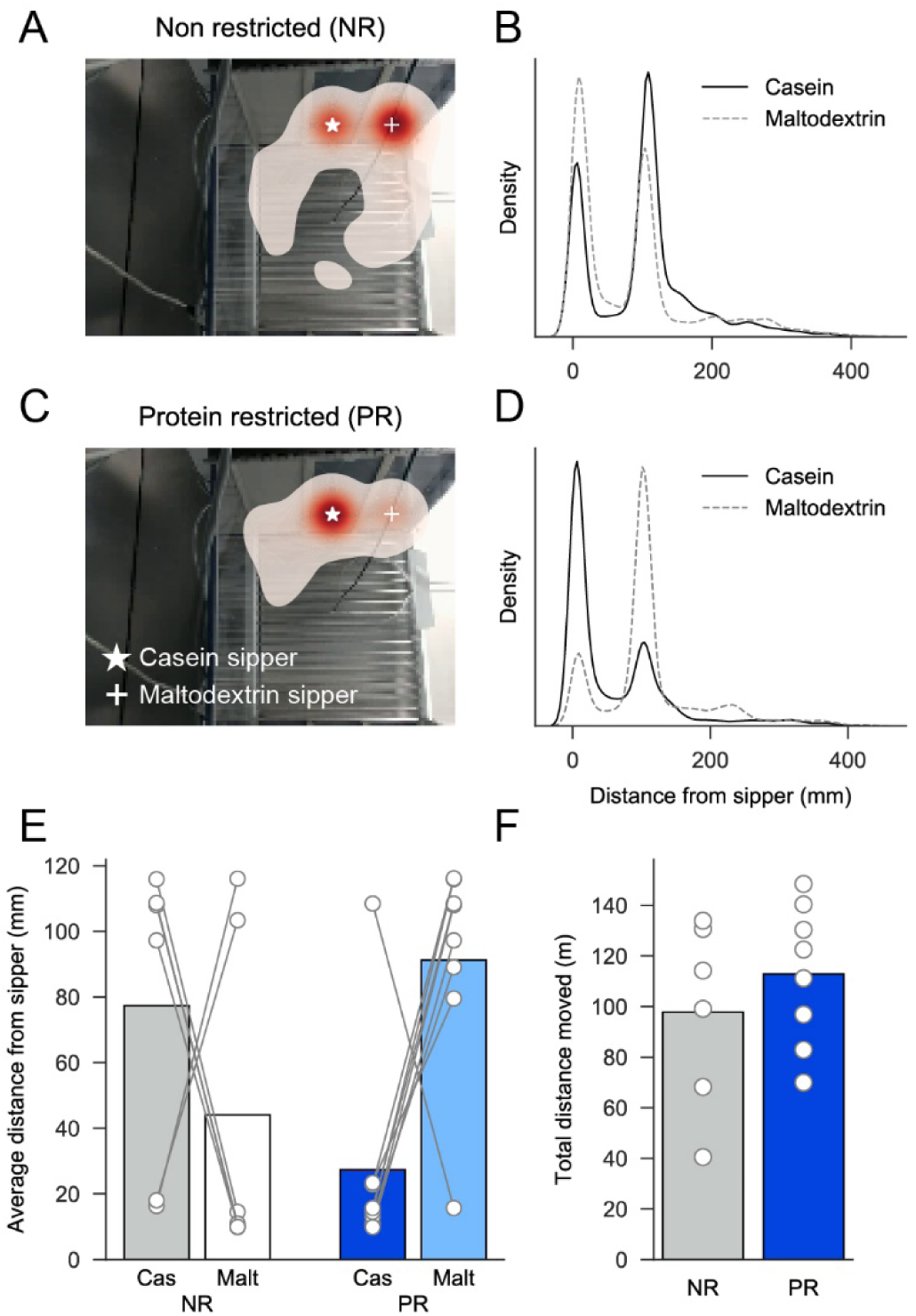
Position in chamber is determined by diet. ***A*,** Heatmap showing position of non-restricted (NR) rat in chamber when tracked across entire session with red colors representing increased time. Casein and maltodextrin sippers are marked with white star and white cross, respectively. ***B*,** Kernel density estimate for all tracked video frames showing distance from casein sipper (black solid line) and maltodextrin sipper (grey dashed line) for position data shown in *A*. ***C-D*,** Same analysis as in *A* and *B* but for a representative protein-restricted (PR) rat. ***E*,** Average distance from each sipper for all rats shows that protein-restricted rats spend more time near the casein sipper than the maltodextrin sipper. Diet x Substance interaction: F(1,12) = 5.03, p=0.045. ***F*,** Total distance moved in entire session is not different between NR and PR rats (t(13)=0.87, p=0.400).

**Extended Data 2-2.**
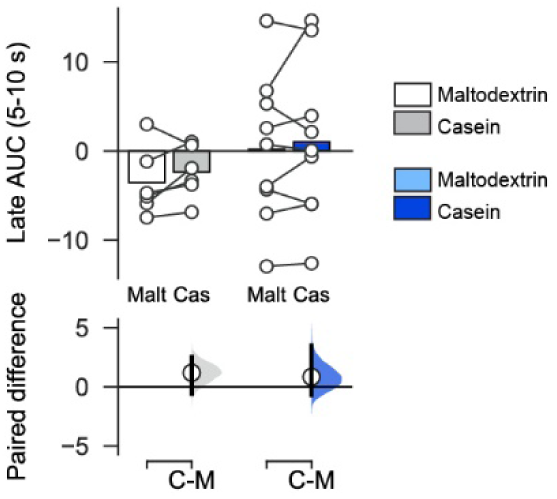
No differences in neural activity identified in period following termination of licking. In the 5 s period following sipper retraction and termination of licking, no difference in the photometry signal is identified due top dietary group or solution (two-way ANOVA, Diet: F(1,13)=0.96, p=0.346; Solution: F(1,13)=1.80, p=0.203; Diet x Solution: F(1,13)=0.05, p=0.824).

**Extended Data 2-3.**
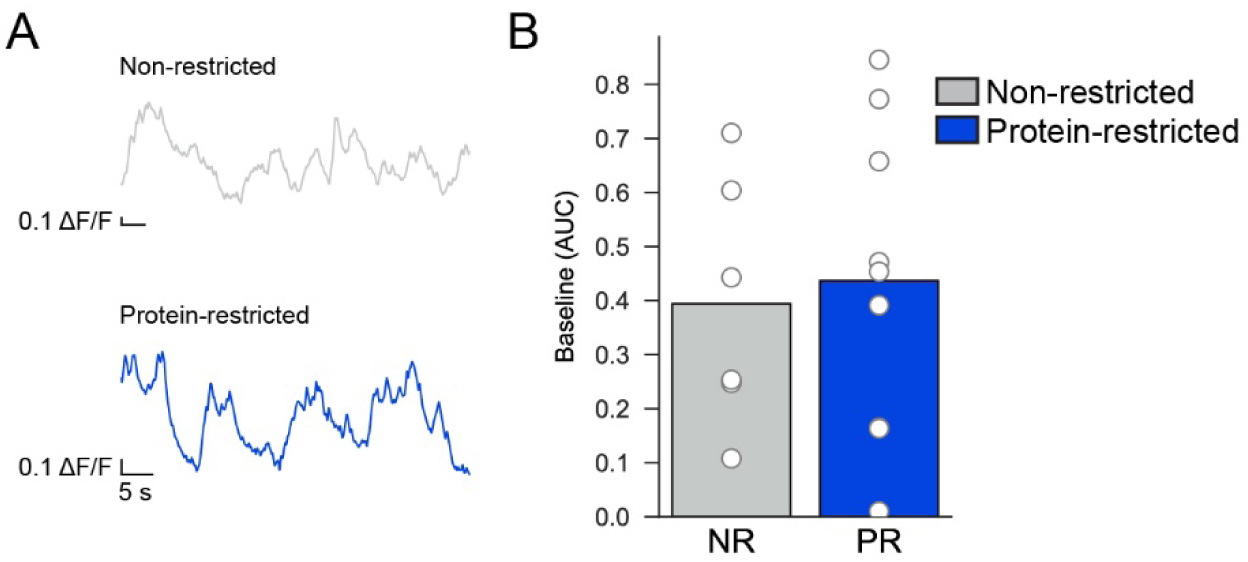
Baseline photometry signal is not affected by diet. ***A,*** Representative traces showing fiber photometry signal at start of session before first sipper extension. Neural activity is observed but not easily quantifiable as distinct transients. ***B,*** Baseline neural activity calculated as AUC of this period is not different between non-restricted (NR) and protein-restricted rats (t(13)=0.30, p=0.769).

**Extended Data 2-4.**
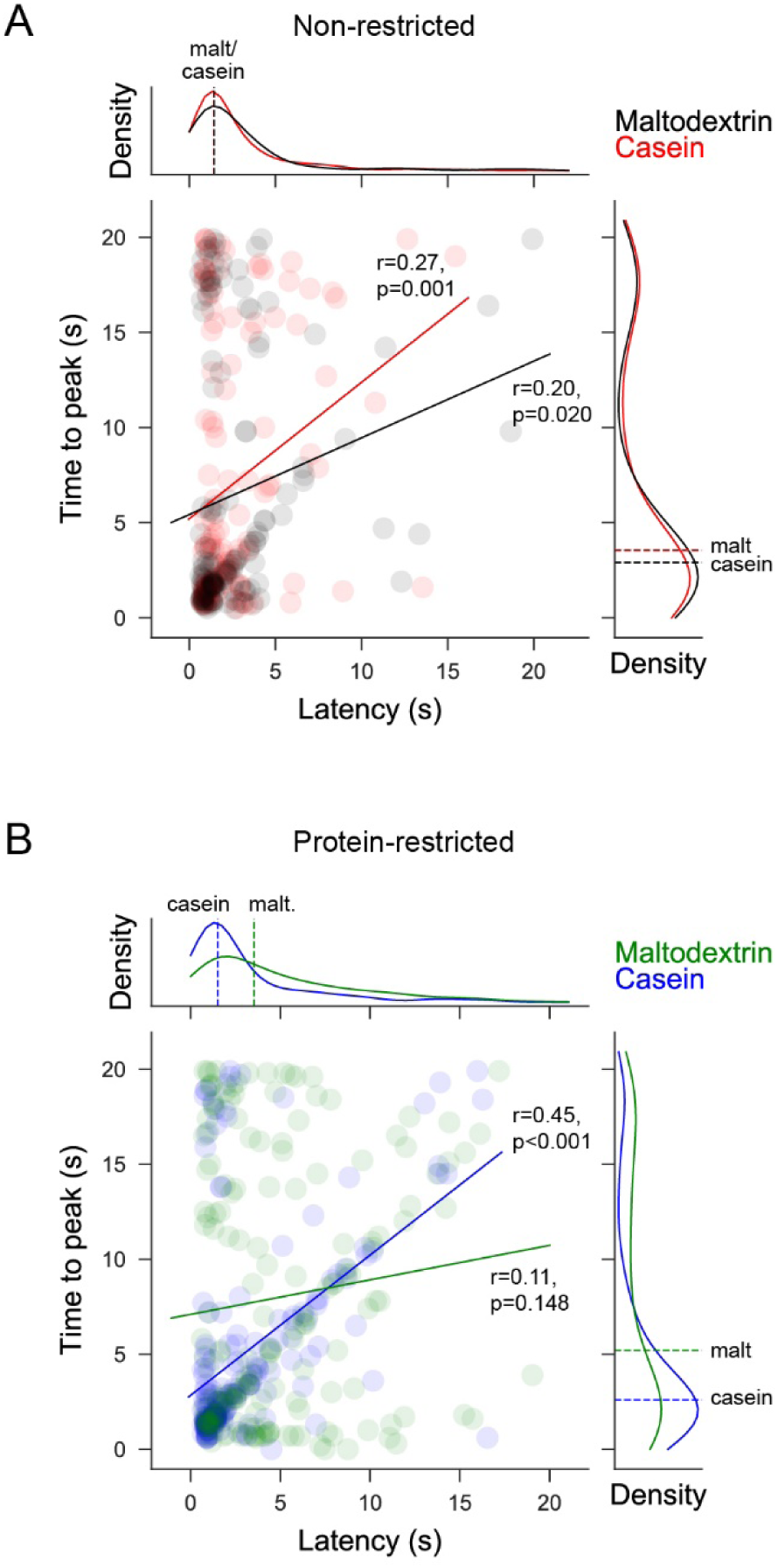
Correlations between latency to start licking following sipper extension and time for photometry signal to peak (from sipper extension) on y-axis. ***A,*** Main plot shows scatter plot of individual trials pooled across all non-restricted rats with latency to lick on x-axis and time for signal to peak on y-axis. Solid line on main plot is linear fit of data with a significant correlation is found for both maltodextrin and casein trials (statistics shown on plot). Density plots are shown for each axis above and to the right, respectively. Dashed lines on density plots show median of data with Mann-Whitney U test showing no difference for either latency to start licking (U=9049, p=0.922) or time to peak (U=9323, p=0.743). Maltodextrin trials are shown in black and casein trials are shown in red. ***B,*** As in A but for protein-restricted rats with maltodextrin trials in green and casein trials in blue. There is a significant correlation for casein trials but not for maltodextrin trials. Comparison of data show that both latency to lick (U=14095, p<0.001) and time to peak (U=16750, p<0.001) are different for maltodextrin and casein trials.

## References

Alhadeff AL, Goldstein N, Park O, Klima ML, Vargas A, Betley JN (2019) Natural and Drug Rewards Engage Distinct Pathways that Converge on Coordinated Hypothalamic and Reward Circuits. Neuron 103:891–908.e6.

Anthony TG, Gietzen DW (2013) Detection of amino acid deprivation in the central nervous system. Curr Opin Clin Nutr Metab Care 16:96–101.

Barbano MF, Cador M (2005) Various aspects of feeding behavior can be partially dissociated in the rat by the incentive properties of food and the physiological state. Behav Neurosci 119:1244–1253.

Bassareo V, Di Chiara G (1997) Differential influence of associative and nonassociative learning mechanisms on the responsiveness of prefrontal and accumbal dopamine transmission to food stimuli in rats fed ad libitum. J Neurosci 17:851–861.

Beeler JA, McCutcheon JE, Cao ZFH, Murakami M, Alexander E, Roitman MF, Zhuang X (2012) Taste uncoupled from nutrition fails to sustain the reinforcing properties of food. Eur J Neurosci 36:2533–2546.

Berridge KC (2007) The debate over dopamine’s role in reward: The case for incentive salience. Psychopharmacology (Berl) 191:391–431.

Berthoud H-R, Münzberg H, Richards BK, Morrison CD (2012) Neural and metabolic regulation of macronutrient intake and selection. Proc Nutr Soc 71:390–400.

Bromberg-Martin ES, Matsumoto M, Hikosaka O (2010) Dopamine in motivational control: rewarding, aversive, and alerting. Neuron 68:815–834.

Brown MTC, Tan KR, O’Connor EC, Nikonenko I, Muller D, Lüscher C, O’Connor EC, Nikonenko I, Muller D, Lüscher C, O’Connor EC, Nikonenko I, Muller D, Lüscher C (2012) Ventral tegmental area GABA projections pause accumbal cholinergic interneurons to enhance associative learning. Nature 492:1–5.

Chaudhari N, Pereira E, Roper SD (2009) Taste receptors for umami: The case for multiple receptors. Am J Clin Nutr 90:738–742.

Cone JJ, Fortin SM, McHenry JA, Stuber GD, McCutcheon JE, Roitman MF (2016) Physiological state gates acquisition and expression of mesolimbic reward prediction signals. Proc Natl Acad Sci U S A 113:1943–1948.

Cone JJ, McCutcheon JE, Roitman MF (2014) Ghrelin Acts as an Interface between Physiological State and Phasic Dopamine Signaling. J Neurosci 34:4905–4913.

Day JJ, Roitman MF, Wightman RM, Carelli RM (2007) Associative learning mediates dynamic shifts in dopamine signaling in the nucleus accumbens. Nat Neurosci 10:1020–1028.

de Araujo IE, Ferreira JG, Tellez LA, Ren X, Yeckel CW (2012) The gut–brain dopamine axis: A regulatory system for caloric intake. Physiol Behav 106:394–399.

de Araujo IE, Oliveira-Maia AJ, Sotnikova TD, Gainetdinov RR, Caron MG, Nicolelis MALL, Simon SA (2008) Food reward in the absence of taste receptor signaling. Neuron 57:930–941.

Di Chiara G, Abizaid A (2009) Ghrelin and dopamine: new insights on the peripheral regulation of appetite. J Neuroendocr 21:787–793.

Dobi A, Margolis EB, Wang H-L, Harvey BK, Morales M (2010) Glutamatergic and nonglutamatergic neurons of the ventral tegmental area establish local synaptic contacts with dopaminergic and nondopaminergic neurons. J Neurosci 30:218–229.

Domingos AI, Vaynshteyn J, Voss HU, Ren X, Gradinaru V, Zang F, Deisseroth K, De Araujo IE, Friedman J (2011) Leptin regulates the reward value of nutrient. Nat Neurosci 14:1562–1568.

Ferreira JG, Tellez LA, Ren X, Yeckel CW, De Araujo IE (2012) Regulation of fat intake in the absence of flavour signalling. J Physiol 590:953–972.

Gietzen DW, Aja SM (2012) The brain’s response to an essential amino acid-deficient diet and the circuitous route to a better meal. Mol Neurobiol 46:332–348.

Godfrey N, Borgland SL (2020) Sex differences in the effect of acute fasting on excitatory and inhibitory synapses onto ventral tegmental area dopamine neurons. J Physiol 598:5523–5539.

Gould JM, Smith PJ, Airey CJ, Mort EJ, Airey LE, Warricker FDM, Pearson-Farr JE, Weston EC, Gould PJW, Semmence OG, Restall KL, Watts JA, McHugh PC, Smith SJ, Dewing JM, Fleming TP, Willaime-Morawek S (2018) Mouse maternal protein restriction during preimplantation alone permanently alters brain neuron proportion and adult short-term memory. Proc Natl Acad Sci 115:E7398–E7407.

Grissom NM, Reyes TM (2013) Gestational overgrowth and undergrowth affect neurodevelopment: Similarities and differences from behavior to epigenetics. Int J Dev Neurosci 31:406–414.

Gunaydin LA, Grosenick L, Finkelstein JC, Kauvar I V, Fenno LE, Adhikari A, Lammel S, Mirzabekov JJ, Airan RD, Zalocusky KA, Tye KM, Anikeeva P, Malenka RC, Deisseroth K (2014) Natural neural projection dynamics underlying social behavior. Cell 157:1535– 1551.

Hajnal A, Smith GP, Norgren R (2004) Oral sucrose stimulation increases accumbens dopamine in the rat. Am J Physiol Regul Integr Comp Physiol 286:R31–7.

Hall KD (2019) The Potential Role of Protein Leverage in the US Obesity Epidemic. Obesity 27:1222–1224.

Heeley N, Blouet C (2016) Central Amino Acid Sensing in the Control of Feeding Behavior. Front Endocrinol (Lausanne) 7.

Heffner TG, Hartman JA, Seiden LS (1980) Feeding increases dopamine metabolism in the rat brain. Science (80-) 208:1168–1170.

Hill CM, Laeger T, Dehner M, Albarado DC, Clarke B, Wanders D, Burke SJ, Collier JJ, Qualls-Creekmore E, Solon-Biet SM, Simpson SJ, Berthoud HR, Münzberg H, Morrison CD (2019) FGF21 Signals Protein Status to the Brain and Adaptively Regulates Food Choice and Metabolism. Cell Rep 27:2934–2947.e3.

Ho J, Tumkaya T, Aryal S, Choi H, Claridge-Chang A (2019) Moving beyond P values: data analysis with estimation graphics. Nat Methods 16:565–566.

Hsu TM, McCutcheon JE, Roitman MF (2018) Parallels and overlap: The integration of homeostatic signals by mesolimbic dopamine neurons. Front Psychiatry 9:1–17.

Ikemoto S, Panksepp J (1999) The role of nucleus accumbens dopamine in motivated behavior: a unifying interpretation with special reference to reward-seeking. Brain Res Brain Res Rev 31:6–41.

Karnani MM, Apergis-Schoute J, Adamantidis A, Jensen LT, de Lecea L, Fugger L, Burdakov D (2011) Activation of central orexin/hypocretin neurons by dietary amino acids. Neuron 72:616–629.

Konanur VR, Hsu TM, Kanoski SE, Hayes MR, Roitman MF (2020) Phasic dopamine responses to a food-predictive cue are suppressed by the glucagon-like peptide-1 receptor agonist Exendin-4. Physiol Behav 215:112771.

Krause EG, Sakai RR (2007) Richter and sodium appetite: from adrenalectomy to molecular biology. Appetite 49:353–367.

Laeger T, Henagan TM, Albarado DC, Redman LM, Bray GA, Noland RC, Münzberg H, Hutson SM, Gettys TW, Schwartz MW, Morrison CD (2014) FGF21 is an endocrine signal of protein restriction. J Clin Invest 124:3913–3922.

Leibowitz SF, Lucas DJ, Leibowitz KL, Jhanwar YS (1991) Developmental patterns of macronutrient intake in female and male rats from weaning to maturity. Physiol Behav 50:1167–1174.

Lerner TNN, Shilyansky C, Davidson TJJ, Evans KEE, Beier KTT, Zalocusky KAA, Crow AKK, Malenka RCC, Luo L, Tomer R, Deisseroth K (2015) Intact-Brain Analyses Reveal Distinct Information Carried by SNc Dopamine Subcircuits. Cell 162:635–647.

Liman ER, Zhang Y V., Montell C (2014) Peripheral coding of taste. Neuron 81:984–1000.

Mathis A, Mamidanna P, Cury KM, Abe T, Murthy VN, Mathis MW, Bethge M (2018) DeepLabCut: markerless pose estimation of user-defined body parts with deep learning. Nat Neurosci 21:1281–1289.

Maurin AC, Jousse C, Averous J, Parry L, Bruhat A, Cherasse Y, Zeng H, Zhang Y, Harding HP, Ron D, Fafournoux P (2005) The GCN2 kinase biases feeding behavior to maintain amino acid homeostasis in omnivores. Cell Metab 1:273–277.

Mayntz D, Raubenheimer D, Salomon M, Toft S, Simpson SJ (2005) Nutrient-specific foraging in invertebrate predators. Science 307:111–113.

McCutcheon JE (2015) The role of dopamine in the pursuit of nutritional value. Physiol Behav 152:408–415.

McCutcheon JE, Beeler JA, Roitman MF (2012a) Sucrose-predictive cues evoke greater phasic dopamine release than saccharin-predictive cues. Synapse 66:346–351.

McCutcheon JE, Ebner SR, Loriaux AL, Roitman MF (2012b) Encoding of aversion by dopamine and the nucleus accumbens. Front Neurosci 6:137.

Mebel DM, Wong JCY, Dong YJ, Borgland SL (2012) Insulin in the ventral tegmental area reduces hedonic feeding and suppresses dopamine concentration via increased reuptake. Eur J Neurosci 36:2336–2346.

Mietlicki-Baase EG, Ortinski PI, Rupprecht LE, Olivos DR, Alhadeff AL, Pierce RC, Hayes MR (2013) The food intake-suppressive effects of glucagon-like peptide-1 receptor signaling in the ventral tegmental area are mediated by AMPA/kainate receptors. Am J Physiol Endocrinol Metab 305:E1367–74.

Mietlicki-Baase EG, Reiner DJ, Cone JJ, Olivos DR, McGrath LE, Zimmer DJ, Roitman MF, Hayes MR (2014) Amylin Modulates the Mesolimbic Dopamine System to Control Energy Balance. Neuropsychopharmacology 40:372–385.

Mohebi A, Pettibone JR, Hamid AA, Wong JMT, Vinson LT, Patriarchi T, Tian L, Kennedy RT, Berke JD (2019) Dissociable dopamine dynamics for learning and motivation. Nature 570:65–70.

Morales M, Margolis EB (2017) Ventral tegmental area: Cellular heterogeneity, connectivity and behaviour. Nat Rev Neurosci 18:73–85.

Morrison CD, Laeger T (2015) Protein-dependent regulation of feeding and metabolism. Trends Endocrinol Metab 26:256–262.

Murphy M, Peters KZ, Denton BS, Lee KA, Chadchankar H, McCutcheon JE (2018) Restriction of dietary protein leads to conditioned protein preference and elevated palatability of protein-containing food in rats. Physiol Behav 184:235–241.

Nair-Roberts RG, Chatelain-Badie SD, Benson E, White-Cooper H, Bolam JP, Ungless MA (2008) Stereological estimates of dopaminergic, GABAergic and glutamatergic neurons in the ventral tegmental area, substantia nigra and retrorubral field in the rat. Neuroscience 152:1024–1031.

Naneix F, Peters KZ, McCutcheon JE (2019) Investigating the effect of physiological need states on palatability and motivation using microstructural analysis of licking Running title : Lick microstructure and physiological states. Neuroscience.

Naneix F, Peters KZ, Young AMJ, McCutcheon JE (2020) Age-dependent effects of protein restriction on dopamine release. Neuropsychopharmacology:1–10.

Nath T, Mathis A, Chen AC, Patel A, Bethge M, Mathis MW (2019) Using DeepLabCut for 3D markerless pose estimation across species and behaviors. Nat Protoc 14:2152– 2176.

Parker NF, Cameron CM, Taliaferro JP, Lee J, Choi JY, Davidson TJ, Daw ND, Witten IB (2016) Reward and choice encoding in terminals of midbrain dopamine neurons depends on striatal target. Nat Neurosci 19:845–854.

Paxinos G, Watson C (1998) The rat brain in stereotaxic coordinates. San Diego: Academic Press.

Phillips PEM, Stuber GD, Heien MLAV, Wightman RM, Carelli RM, Hill C (2003) Subsecond dopamine release promotes cocaine seeking. Nature 422:614–618.

Raubenheimer D, Simpson SJ (2019) Protein Leverage: Theoretical Foundations and Ten Points of Clarification. Obesity 27:1225–1238.

Robinson MJF, Berridge KC (2013) Instant transformation of learned repulsion into motivational “wanting”. Curr Biol 23:282–289.

Roitman MF, Wheeler RA, Wightman RM, Carelli RM (2008) Real-time chemical responses in the nucleus accumbens differentiate rewarding and aversive stimuli. Nat Neurosci 11:1376–1377.

Root DH, Barker DJ, Estrin DJ, Miranda-Barrientos JA, Liu B, Zhang S, Wang HL, Vautier F, Ramakrishnan C, Kim YS, Fenno L, Deisseroth K, Morales M (2020) Distinct Signaling by Ventral Tegmental Area Glutamate, GABA, and Combinatorial Glutamate-GABA Neurons in Motivated Behavior. Cell Rep 32:108094.

Salamone JD, Correa M (2012) The mysterious motivational functions of mesolimbic dopamine. Neuron 76:470–485.

Sclafani A, Touzani K, Bodnar RJ (2011) Dopamine and learned food preferences. Physiol Behav 104:64–68.

Simpson SJ, Raubenheimer D (2005) Obesity: The protein leverage hypothesis. Obes Rev 6:133–142.

Sulzer D, Cragg SJ, Rice ME (2016) Striatal dopamine neurotransmission: Regulation of release and uptake. Elsevier GmbH.

Tellez LA, Han W, Zhang X, Ferreira TL, Perez IO, Shammah-Lagnado SJ, van den Pol AN, de Araujo IE (2016) Separate circuitries encode the hedonic and nutritional values of sugar. Nat Neurosci 19:465–470.

Theall CL, Wurtman JJ, Wurtman RJ (1984) Self-selection and regulation of protein: carbohydrate ratio in foods adult rats eat. J Nutr 114:711–718.

Tsang AH, Nuzzaci D, Darwish T, Samudrala H, Blouet C (2020) Nutrient sensing in the nucleus of the solitary tract mediates non-aversive suppression of feeding via inhibition of AgRP neurons. Mol Metab 42:101070.

Wurtman JJ, Baum MJ (1980) Estrogen reduces total food and carbohydrate intake, but not protein intake, in female rats. Physiol Behav 24:823–827.

Zessen R Van, Phillips JL, Budygin EA, Stuber GD (2012) Article Activation of VTA GABA Neurons Disrupts Reward Consumption. Neuron 73:1184–1194.

